# Actin-binding compounds that affect the kinetics of the interaction of cardiac myosin with actin

**DOI:** 10.1101/2020.09.16.300517

**Authors:** Osha Roopnarine, David D. Thomas

## Abstract

We measured the effects of ten actin-binding compounds on the interaction of cardiac myosin subfragment 1 (S1) with pyrene labeled F-actin (PFA). These compounds, previously identified from a small-molecule high-throughput screen (HTS), perturb the microsecond structural dynamics of actin and the steady-state activity of actin-activated myosin ATPase. In the present study, we have further characterized their mechanisms of action by measuring their effects on PFA fluorescence, which is decreased specifically by the strong binding of myosin to actin, and is restored upon release of S1 by MgATP. We measured the effects of compounds under equilibrium and steady-state conditions, as affected by S1 and ATP, and also under transient conditions, in stopped-flow experiments following rapid addition of ATP to S1-bound PFA. We observe that these compounds affect the early steps of the myosin ATPase cycle to different extents (mild, moderate, and severe). The compounds decrease the equilibrium constant for the formation of the collision complex and the rate constant for subsequent isomerization to the ternary complex, indicating increased ATP affinity and trapping of ATP in the myosin active site. These compound effects on actin structure inhibit the kinetics of the actin-myosin interaction in ways that may be desirable for possible treatment of hypercontractile forms of hypertrophic cardiomyopathy (HCM). This work helps to elucidate the mechanisms of action of these compounds, several of which are currently used therapeutically, and it sets the stage for future HTS campaigns on a larger scale, to discover new drugs for treatment of heart failure.

## Introduction

Actin is abundant in eukaryotic cells and is involved in many important cellular processes such as movement, cell division, maintenance of cellular shape, transport of vesicles, phagocytosis, and contractility (1). Actin interacts with and is regulated by many actin binding proteins, such as capping and severing proteins, nucleating proteins, stabilizers, and contractile proteins such as tropomyosin (TM) and troponin complex (Tn) to form thin filaments, myosin, and myosin binding protein C.

In striated muscle contraction, actin and myosin interactions are nucleotide-dependent, as the hydrolysis of ATP provides energy for the conformational and structural changes that result in force. In rigor, actin (A) and myosin (M) binds strongly (S) in a stereospecific manner in the absence of ATP to form the actin-myosin complex (A.M), but this changes upon ATP (T) binding to the myosin active site (A.M.T), which induces weak-binding (W) and rapid dissociation of myosin from actin (M.ATP). Then hydrolysis of the bound ATP (M.ADP.P_i_) induces conformational changes in myosin that reorient the lever arm (consisting of the converter region, and the light chain binding region that contains the essential and regulatory light chains) to prime myosin for weak binding (W) reattachment to actin (A.M.ADP.P_i_) that is followed by the release of P_i_, reorientation of the lever arm, and the powerstroke (force producing) to a strongly-bound (S) A.M.ADP state. In cardiac muscle contraction, the release of ADP to form the A.M state is a rate-limiting step that completes the myosin ATPase cycle (2). Therefore, a key actin-myosin interaction during force production is the weak-to-strong (W→S) transition that myosin undergoes in the ATPase cycle (3) that can be regulated by several structural and kinetic steps (2).

Actin has clinical relevance, as both skeletal and cardiac actin isoforms have mutations that cause human diseases. Skeletal muscle actin mutations account for about 20% of congenital myopathies and occur as nemaline, intranuclear rod and/or actin myopathy, and congenital fiber type disproportion (4). Even more importantly, cardiac actin mutations that appear in the myosin or tropomyosin binding regions cause either hypertrophic or dilated cardiomyopathies (5), increasing the motivation to discover small molecules that target actin for therapeutic treatment of actin-related diseases.

Small molecules have recently been investigated as potential therapeutic agents for myosin mutations that cause inherited heart disease, with particular focus on hypertrophic cardiomyopathy (HCM). These potential drugs include omecamtiv mecarbil (OM) (6,7) and mavacamten (Mava), formerly known as MYK 461 (8), both of which were developed from HTS screens targeting myosin. It was shown that OM increases contractility of cardiac muscle and is an activator of actin-activated myosin ATPase (6) (7,9). More recently, OM was shown to activate the cardiac ryanodine receptor (RyR) calcium release channel, which may promote a pro-arrhythmic risk (10,11). Mava is in the third phase of clinical trials as a therapeutic for hypercontractility in HCM caused by myosin mutations (12). Recently, Mava was also proposed for treatment of HCM caused by mutations in the calcium regulatory thin filament protein troponin (13). However, the results of that study were mixed, suggesting that for these troponin-based HCM mutations, Mava would not be a suitable therapeutic candidate.

Small-molecule compounds designed for use as drugs to manage specific human diseases may affect other systems in the body, including the cardiovascular system. Therefore, it is important to determine the effects of these compounds on the contractility of cardiac and skeletal muscle. Because actin’s role in muscle contractility (both striated and smooth) is regulated by many other proteins such as in the thin filament proteins (tropomyosin, troponin, caldesmon, etc) and in thick filament proteins (myosin heavy chain, myosin light chains, myosin binding protein C), it is also important to determine the effects of the actin-binding compounds on these associated proteins’ function. The focus of the present study is to determine how previously identified actin-binding compounds affect cardiac myosin’s ATPase function.

A recently engineered fluorescence resonance energy transfer (FRET) biosensor, consisting of donor-labeled actin and an acceptor-labeled peptide derived from the N-terminus of the myosin essential light chain (ELC), was used in a high-throughput screen (HTS) of the NIH Clinical Collection (NCC), a library of compounds that have a history of use and safety in human clinical trials (14,15). In that study, ten “hit” compounds (Figure 1) from NCC were found to affect actin-myosin function. These ten compounds are FDA-approved drugs for therapeutic treatment in diverse human diseases (Figure 1). Car is used as a β-blocker for treatment of hypertension, Dan is used as a skeletal muscle relaxant, and Mef is used as an anti-malarial. Thi, Flu, and Phn are used as anti-psychotics. Flp, Hon, and Mit are used to treat cancer. Teg has been used to treat irritable bowel syndrome and constipation. However, Thi and Flp are no longer in common use because they cause liver toxicity, Mit is under FDA caution as it may contribute to heart failure, and Teg is no longer in use because it can contribute to heart failure. All ten of these compounds significantly decreased the FRET efficiency in the above-described high-throughput screen, indicating decreased myosin-actin interaction. Some of the compounds also had significant effects on actin filament flexibility, and on steady-state actin-activated myosin S1 ATPase activities (14,15).

**Figure 1.**
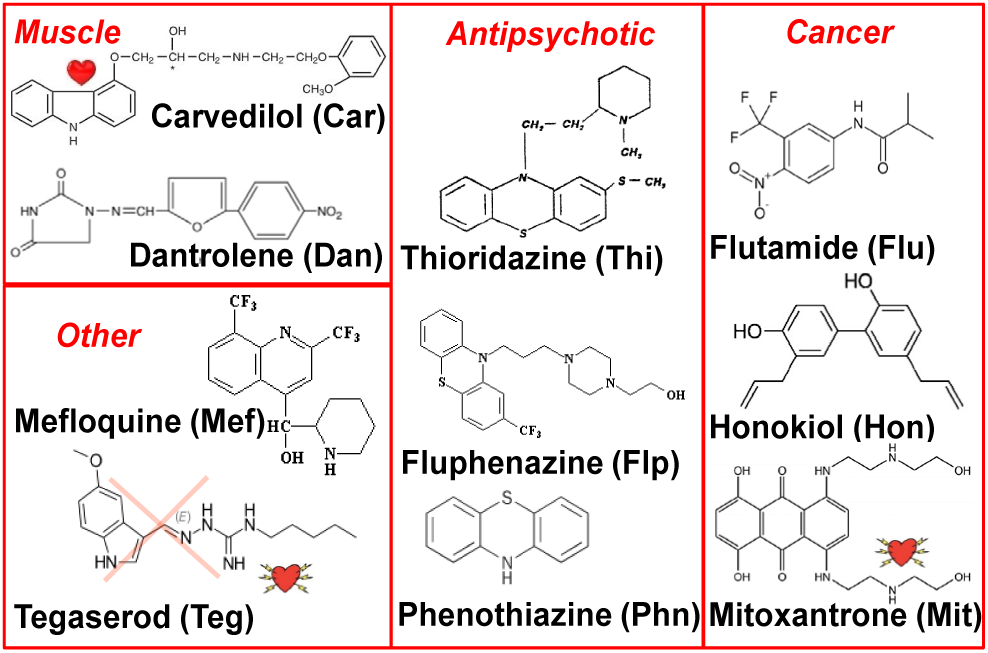
Chemical structures for ten actin-binding compounds discovered from a HTS FRET assay. The drugs are categorized by their intended application for treatment of human disease (see text for usage and warnings). The drugs will be referred to below by the indicated abbreviations.

A subsequent study by the same authors showed that some of these actin-binding compounds have distinct effects on the polymerization properties of skeletal and cardiac actin isoforms, and on the steady-state ATPase activities of skeletal and cardiac myofibrillar ATPase activity (16). Given the very small sequence differences in structure between cardiac and skeletal muscle actin, this is a surprising finding. This is also important, because it establishes feasibility for development of actin-targeted drugs for specific muscle types.

In the present study, we investigate in more detail the mechanisms of action for these actin-binding compounds, focusing on cardiac muscle because of its high potential therapeutic significance. Specifically, we ask the following questions: Which steps in the actin-myosin ATPase reaction do these compounds affect? What are the effects of these compounds on the *rates of transition between weak and strong actin-myosin binding?* A key tool in answering these questions is the fluorescence of pyrene-labeled actin, which has been well characterized for detecting the kinetics of association and disassociation of strong-binding myosin heads on actin (17).

Pyrene iodoacetamide (PIA) covalently binds to cysteine 374 (C374) in F-actin, which is proximal to, but not within the myosin interaction sites on actin. C374 does not appear to be directly involved in the myosin-actin interaction (18), but rather it might be important for monomer-to-monomer contact in the actin filament, because deletion of C374 destabilizes F-actin (19). Labeling of C374 with PIA, producing pyrene-labeled F-actin (PFA), has no significant effects on the equilibrium and dynamics of myosin interaction with actin, but when S1 binds to PFA, the fluorescence is decreased because a local conformational change causes the pyrene moiety to become more accessible to the solution (20,21). The decrease of PFA fluorescence in the presence of a compound indicates that the compound binds to actin and induces a local conformational change that exposes the pyrene moiety to the solvent.

In the present study, we labeled F-actin with pyrene-iodoacetamide to detect the impact of the compounds on the steady-state PFA fluorescence in the absence and presence of bovine cardiac ventricular myosin S1 (β-S1) and ATP, in order to determine the effects of compounds on the transition from the weak-to strong-binding states. We then used stopped-flow fluorescence to study the transient-state kinetics of the formation of the actin-S1 complex and the effects of ATP (T) binding to the myosin (M) active site in the presence and absence of the compounds. Specifically, we determined how the compounds affect the binding affinity of actin and S1 in rigor, the ATP affinity for myosin in the myosin-actin complex, and the subsequent rate of isomerization of the collision complex (A.M.T) to the ternary complex (A∼M.T). We found that most of these compounds weaken the actin-myosin interaction in rigor, increase the ATP affinity for myosin, and decrease the rate constant of the subsequent step in the ATPase cycle. These results support the hypothesis that the compounds inhibit the actin-myosin interaction through the mechanism of trapping ATP in the myosin active site.

## Results

### Characterization of pyrene-labeled actin

The ratio of bound pyrene to F-actin was 0.95-1.1, consistent with complete and specific reaction at Cys374 (17). Bovine ventricular myosin S1 (0.1 μM) had Mg-ATPase activity of 0.009 ± 0.04 s^-1^ in the absence of actin and 1.15 ± 0.03 s^-1^ in the presence of 15 μM phalloidin-stabilized F-actin, consistent with previous results (22,23).

### Pyrene-Actin detects the weak-to-strong transition in the actin-myosin interaction

As shown previously (17,20), phalloidin-stabilized pyrene-labeled F-actin (PFA) has high steady-state fluorescence intensity with a maximum at 407 nm (Figure 2A, black spectrum), and the addition of myosin S1 to pyrene-labeled F-actin linearly decreases the fluorescence intensity (Figure 2A, green spectrum) until the molar ratio of S1 to Actin is 1, and then it becomes independent of [S1] (Figure 2B). When saturating MgATP binds to myosin’s active site, it weakens the interaction of myosin S1 with actin, completely restoring pyrene fluorescence (Figure 2 red).

**Figure 2.**
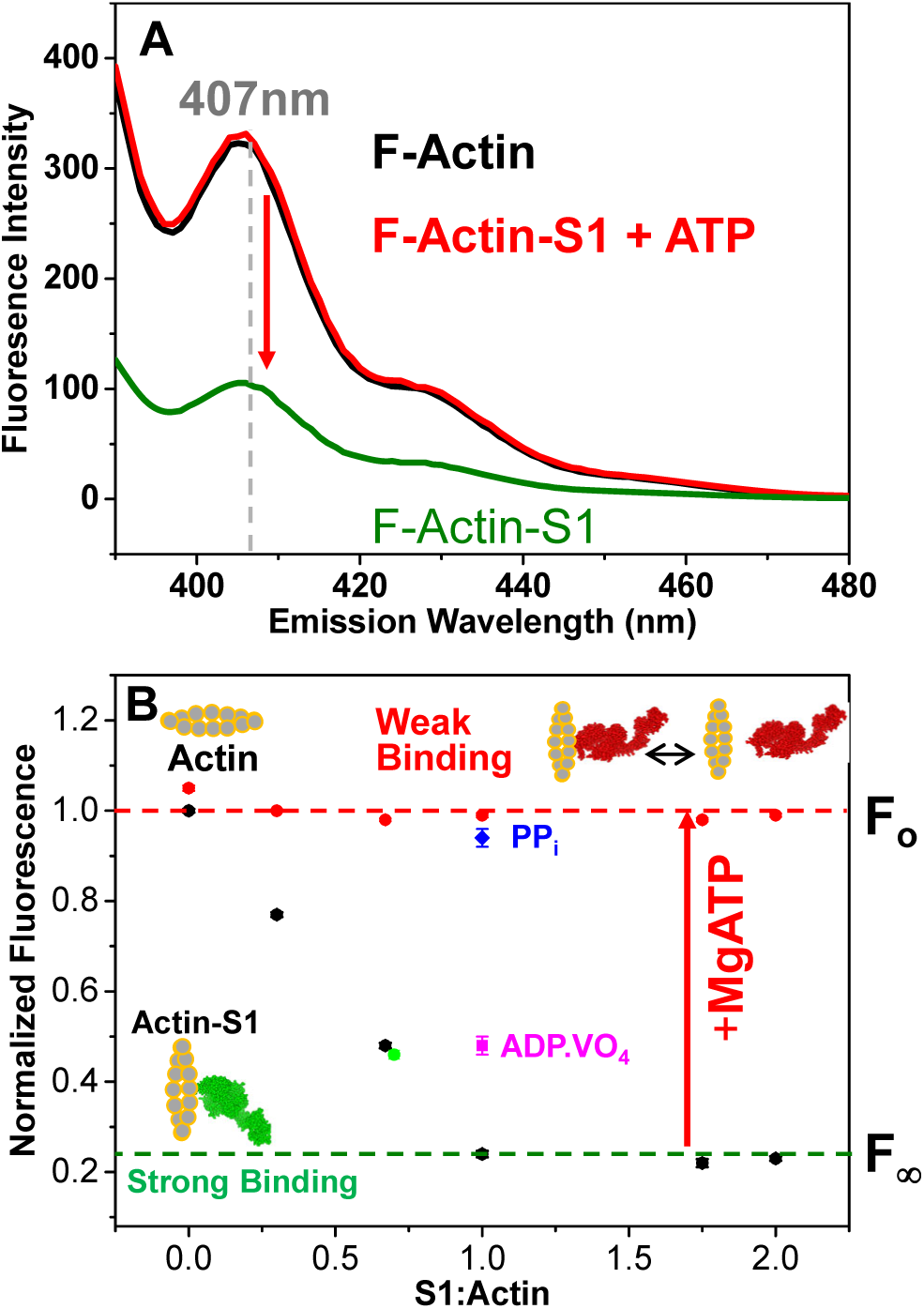
**A**: Steady-state fluorescence spectra of 2μM PFA (black) as a function of [S1] in the absence (green) and presence (red) of Mg.ATP. **B**: Inhibition of PFA fluorescence as a function of [S1] (•) and restoration of PFA fluorescence by MgATP (•). Addition of 3mM ADP.VO_4_ (▪) and 10mM PP_i_ (♦) to the PFA-S1 complex.0.7:1 S1:Actin is shown in green (•).

The formation of the ADP.VO_4_-S1 complex relieved the S1-induced inhibition of the pyrene fluorescence to 48 ± 3%, showing that PFA is sensitive to intermediate states of the ATPase cycle (Figure 2B (▪)). Vanadate is a phosphate analog that, together with ADP (when vanadate is added during cycling ATP), traps myosin heads in a weak-binding state similar to the M.ADP.P_i_ state (24,25). The addition of 10 mM potassium pyrophosphate (PP_i_) to the PFA-S1 almost completely relieved the inhibition of the pyrene fluorescence (Figure 2B, (♦)). Pyrophosphate is known to dissociate the rigor actin-myosin complex similarly to MgATP (26,27).

### Controls with myosin-specific binding compounds: OM and Mava

Omecamtiv mecarbil (OM) and mavacamten (Mava), which bind specifically to myosin (8), were used as controls to demonstrate that changes in pyrene fluorescence are due to changes in actin-myosin interaction. The addition of 5 μM OM or Mava did not induce significant changes in the PFA fluorescence, (4% and 3%) (Figure 3, left, purple and pink bars). When S1 was added (equimolar to actin), the expected decrease in pyrene fluorescence was observed (Figure 3, middle, green), with no effect of OM or Mava (Figure 3, middle, purple and pink bars). Addition of MgATP partially restored PFA fluorescence, to 95% in the absence of compound, and to 65% in the presence of OM and to 84% in the presence of Mava (Figure 3, suggesting that these compounds partially stabilize the strong-binding state of myosin and actin.

**Figure 3.**
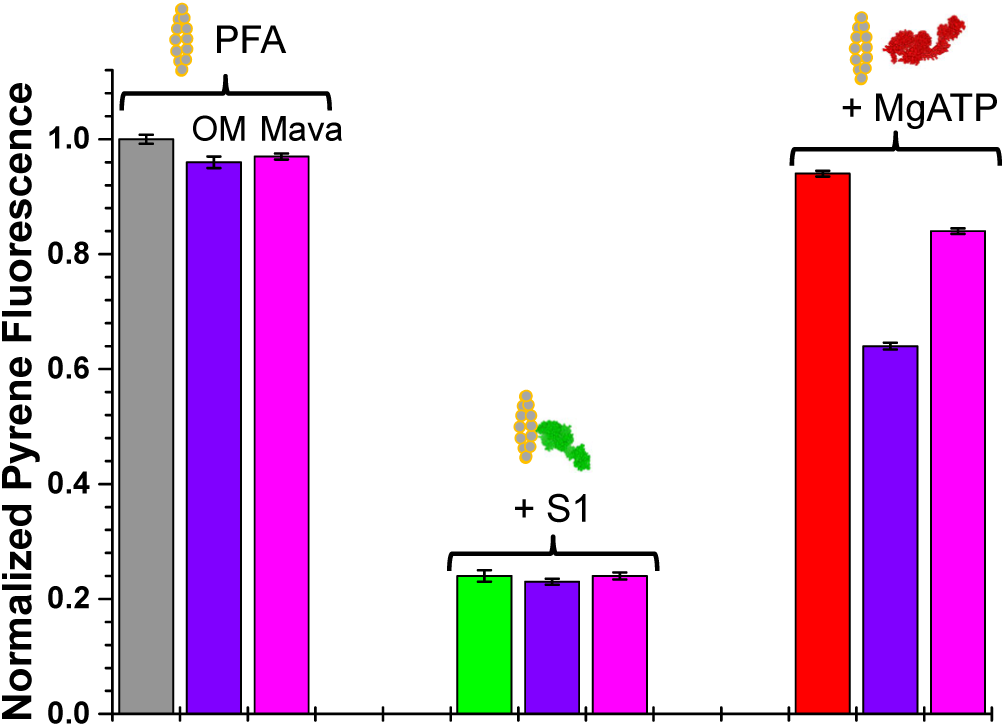
Effect of myosin-binding drugs, OM (purple) and Mava (pink) on the steady-state fluorescence of 2 μM PFA (left, grey), 2 μM PFA.S1 (green) in the absence (middle) and in presence (red, right) of MgATP.

### Effects of compounds on PFA fluorescence

Control experiments with the compounds (100 μM) showed minimal emission at 407 nm (0-1.6%), suggesting that the compounds do not contribute to PFA fluorescence emission at 407 nm. The dose-response curves of the compounds in the range of 2.5-150 μM compound show that Car, Teg, Phe, and Hon have minimal effects on the pyrene fluorescence, therefore the K_d_ was not determined for these compounds. However, Mef, Flp, Thi, Flu, Dan and Mit decreased PFA fluorescence with hyperbolic dependence (Figure 4A, B), so the apparent dissociation equilibrium constant, K_d_, was determined from fits to a hyperbolic equation (Experimental procedures: Eq. 2 and 3), as shown for six compounds (Figure 4A, C). Mit, Flup, Dan, and Mef had the smallest K_d_ (5.2 ± 1.2 μM, 7.7 ± 2.8 μM, 16.3 ± 6 μM, 20 ± 10 µM, respectively), indicating strong binding to actin. Thi and Flu had K_d_ values of 38.8 ± 10 μM and 44.8 ± 22 μM respectively. MgATP had minimal effect on the fluorescence of PFA-Compound complexes, except for Flp, Mef, and Mit, which partially reversed the decrease of PFA fluorescence. The ATP-induced alleviation of fluorescence decrease with Mit required higher concentrations of MgATP (up to 400 mM) to relieve the inhibition (Figure 4D), where the K_d_ increased from 5.2 ± 1.2 µM to 24.7 ± 5.8 μM, suggesting weaker binding of Mit in the presence of excess ATP. Similarly, higher [ATP] also increased the K_d_ of Flp to 25.8 ± 14.2 μM and Mef to 89 ± 28.4 μM.

**Figure 4.**
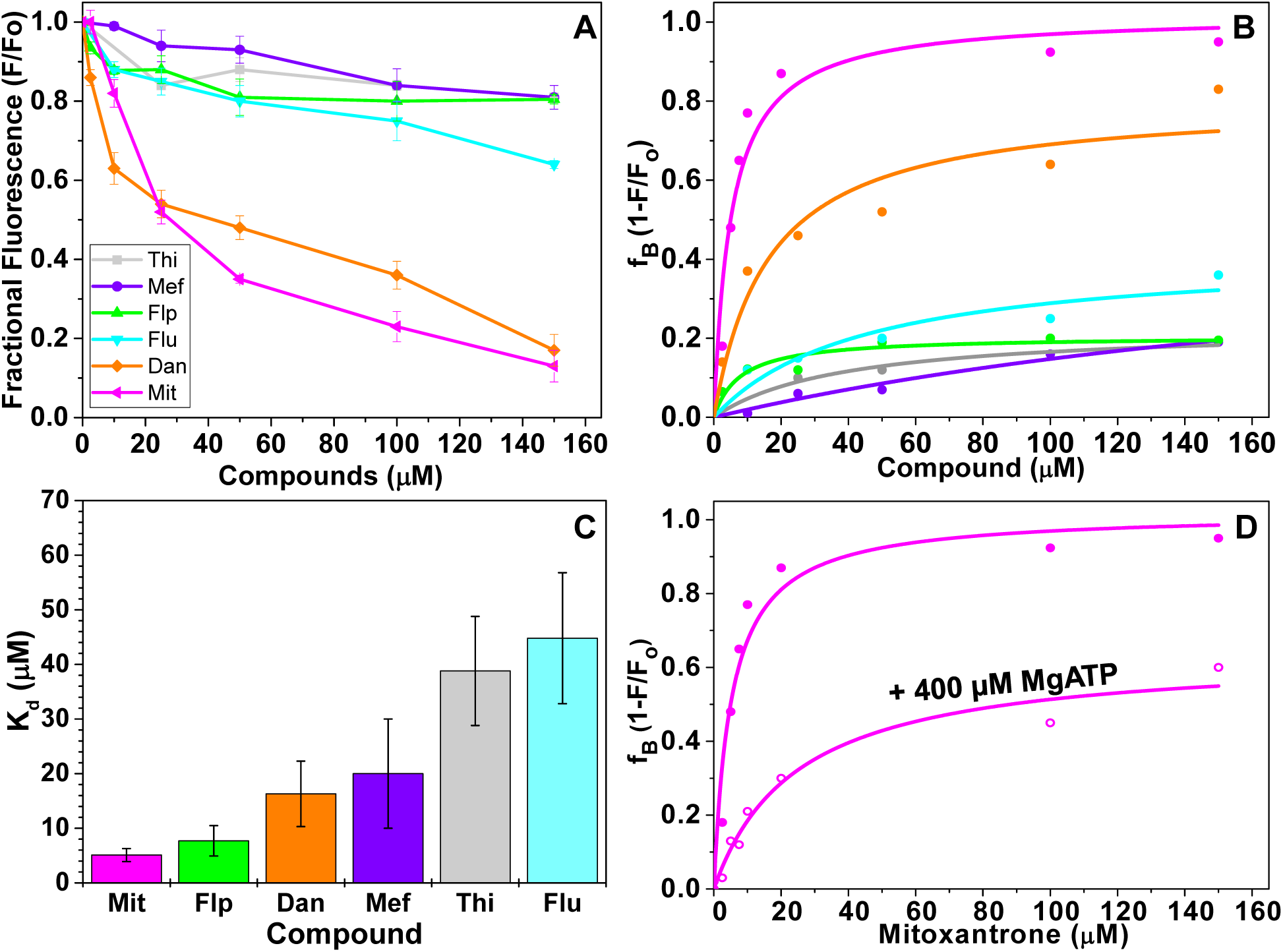
Effects of compounds on the steady-state fluorescence of PFA (2μM). **A**: Dose response of compounds on PFA; **B**: the hyperbolic fits of the fraction bound (f_b_) vs [compound] for some of the compounds; **C**: the dissociation equilibrium constant, K_d_; **D**: the hyperbolic dependence of Mit in the absence (•) and in the presence (○) of MgATP.

The addition of 100 μM compounds to 2 μM PFA decreased the steady-state pyrene fluorescence to varying extents (Figure 5), indicating that these compounds affect the structure of actin in the C-terminal region, as previously observed with other optical probes attached to C374 (14). The following experiments were conducted using [compound] that was at least twice the K_d_ value: 100 μM compound except for Mit (10 μM), Flu and Dan (50 μM).

**Figure 5.**
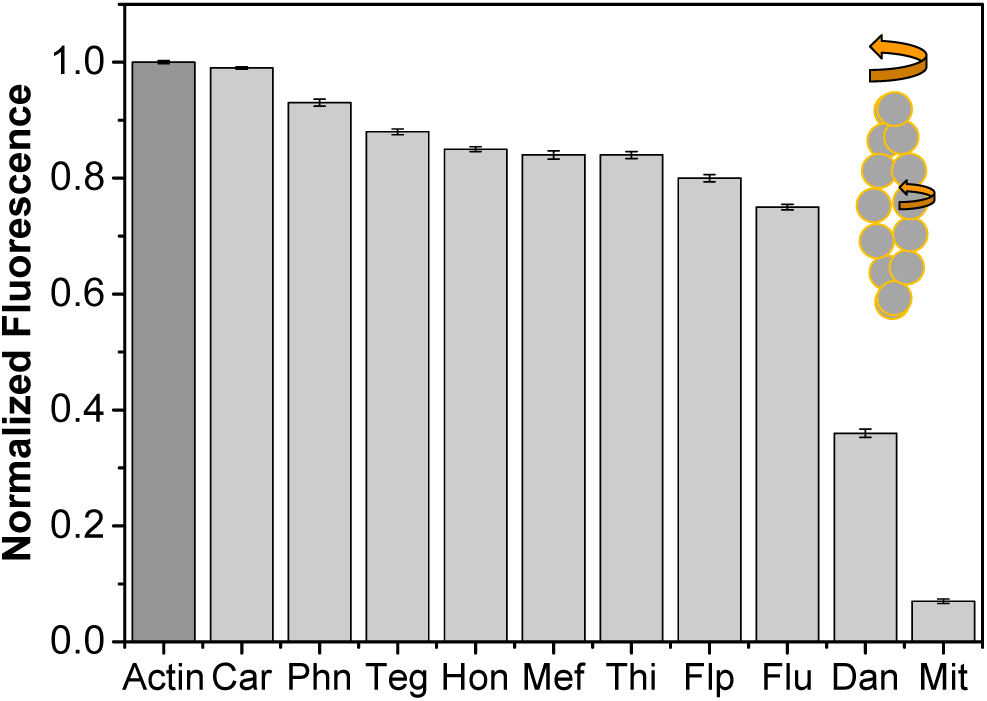
Normalized fluorescence of PFA in the absence (dark grey) and in the presence (grey) of 100 μM compound. Inset: possible conformational changes in actin due to the compound.

### Effects of the compounds on S1 binding and dissociation from F-Actin

To determine the effect of each compound (C) on S1 binding and ATP-induced dissociation from actin, the strategy was to first create the PFA.C complex and measure PFA fluorescence (Figure 6, grey bar), then create the PFA.C.S1 complex by adding a substoichiometric ratio amount (0.7:1) of cardiac S1 and measure PFA fluorescence again (shown in Figure 6A, pink bar). We then added Mg.ATP and measured PFA fluorescence (Figure 6B, peach bars). As shown in Figure 2B (green circle), the addition of cardiac S1, at a molar ratio of 0.7:1 with actin, induced a 46 ± 0.1% decrease in pyrene fluorescence intensity (Figure 6, “S1”). Comparison of the calculated PFA fluorescence (*F*_calc_, Eq. 4) (Figure 6A, hatched bars) and experimental values (Figure 6A, green bars) of the compound and S1, shows that some of the compounds (Teg, Flp, Flu, Mef, Mit, and Dan) induced a smaller decrease in PFA fluorescence than expected, suggesting the presence of weak-binding myosin. Other compounds (Car, Thi, and Phe) induced a greater decrease in PFA fluorescence, suggesting stronger binding of S1 to actin in the presence of these compounds, and Hon had no effect. The addition of ATP to the PFA-S1 rigor complex (Figure 6B, peach bars), in the absence of the compound, dissociated the bound S1 and induced almost full recovery of the PFA fluorescence (Figure 6B, pink). However, in the presence of several compounds, the addition of ATP to the PFA.C.S1 rigor complex prevented full restoration of PFA fluorescence (except for Car), suggesting that the ATP-induced release of S1 was perturbed in the presence of the compound (Figure 6B, peach bars).

**Figure 6.**
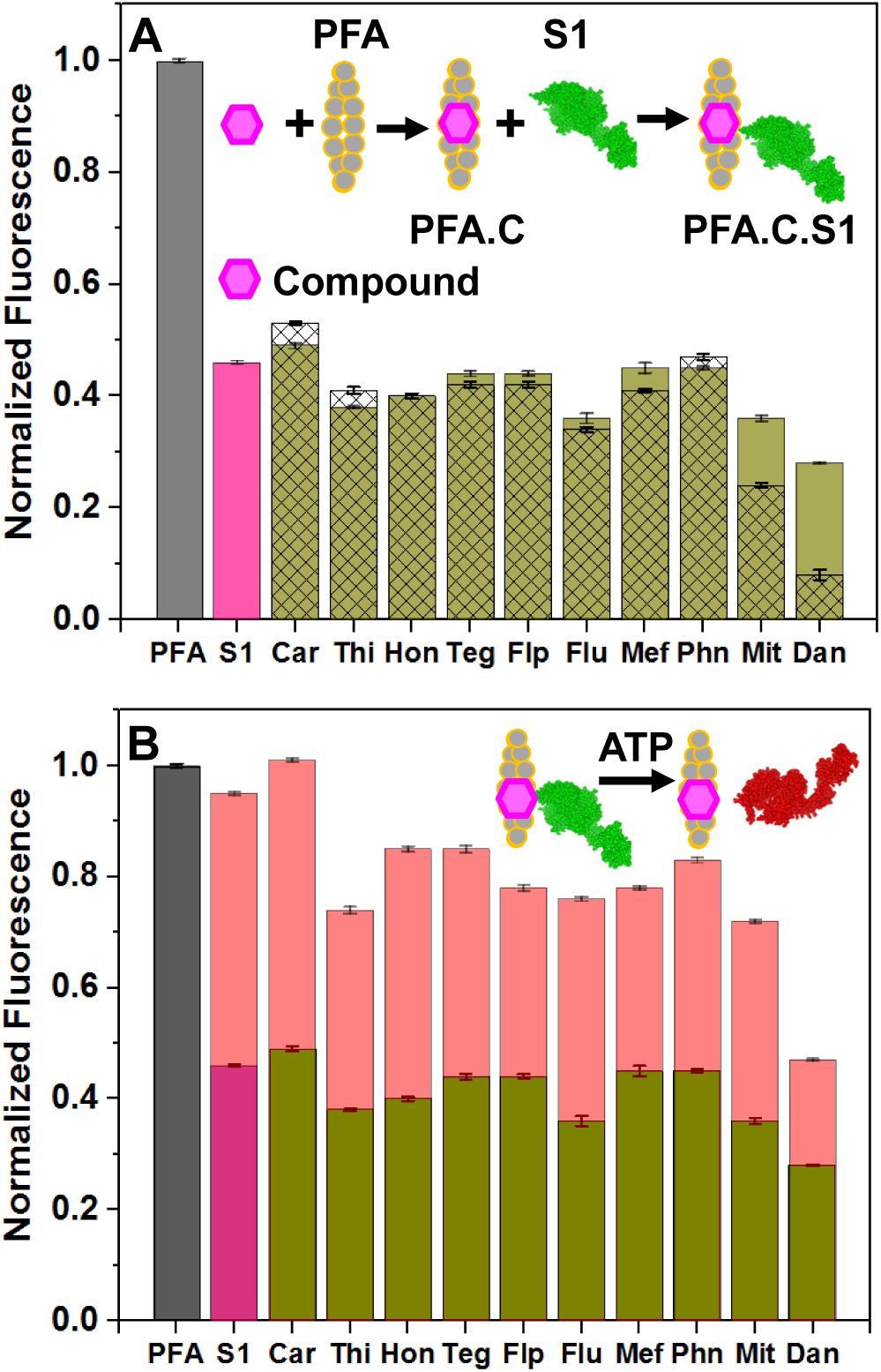
Normalized steady-state fluorescence of 2μM PFA (grey) with S1 (0.7 S1/Actin) only (pink). **A**: Calculated (hatched bars) and experimental (green) normalized fluorescence of PFA.Compound.S1 complex. **B**: The effect of ATP (peach) on the PFA.Compound.S1 complex (green).

### Transient State Kinetics

#### Rigor binding of actin-compound complex to S1

The rate of binding of actin and S1, *k*_+A_, was determined by varying the PFA concentration (0.04-0.24 μM) in the presence of S1 (0.02 μM) in the absence (control) and presence of the compound (used in the same molar ratio to actin as in the steady-state fluorescence experiments; PFA:compound for Mit was 1:5, and for Dan and Flu 1:25). The fluorescence of the PFA in the presence of S1 decreased as a function of time and was fitted to a single exponential (Figure 7A). The experiments were done in the presence of apyrase, which removed the fast component that may be due to any residual ATP (See Experimental procedures: **Rigor binding of S1 to Actin**) (17,28). The rate constant, *k*_obs_, was linearly dependent on [actin], and the slope of the plot of *k*_obs_ vs [actin] yielded the second-order binding constant, *k*_+A,_ of actin and S1, while the intercept gave the rate of dissociation, *k*_-A_. The dissociation equilibrium constant, *K*_A_, was determined from *k*_-A_/*k*_+A_.

**Figure 7.**
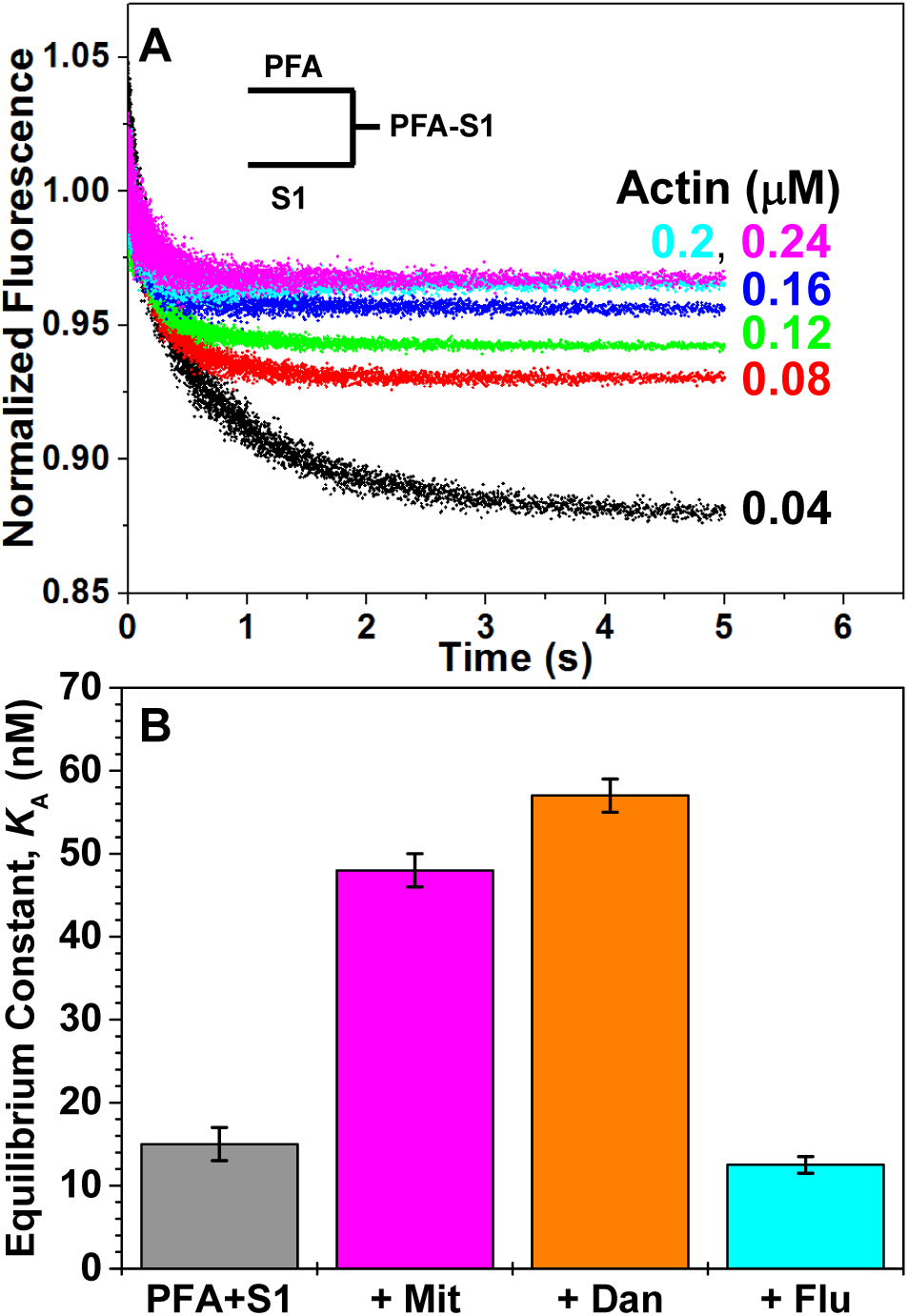
**A**: Normalized fluorescence transients of varying [PFA] with β-S1 in the absence of compounds. **B**: Equilibrium constants for PFA + β-S1 in the presence of Mit, Dan, and Flu.

Despite the addition of apyrase to remove any contaminating nucleotide, Dan and Mit induced second component with a slower rate, *k*_obs_,_slow_, that was insensitive to the [actin]. For PFA binding to β-S1, the equilibrium constant, *K*_A_, was 15 ± 1 nM, which increased in the presence of Mit and Dan, to 48 ± 2 nM and 57 ± 2 nM respectively, suggesting that these compounds decreased the affinity of actin binding to S1 more significantly than Flu, which induced a dissociation equilibrium constant of 13 ± 1 nM (Figure 7B). *K*_A_ was not determined in the presence of the other compounds.

### ATP-induced release of S1 from actin-compound-S1 complex

To determine the equilibrium constant, 1/*K*_1_, for the formation of the collision complex, A.M.T, the rigor complex of pyrene-labeled-actin (PFA) and myosin-S1 complex was formed in one syringe and the Mg.ATP placed in another syringe. Rapid mixing results in a rapid increase of pyrene fluorescence as ATP dissociates the S1 (Figure 8A). This biphasic transient curve is best fitted to a double exponential equation to yield a fast rate constant *k*_obs1_ and slow *k*_obs2_, with corresponding large amplitude, *A*_1_ and small amplitude *A*_2_. For the transient curve in Figure 8A, 50 nM PFA and S1 (1:1) in 25 μM MgATP (final concentrations) has a fast phase and large amplitude (118 ± 7 s^-1^, 80%) and slow phase with a small amplitude (20 ± 3.5 s^-1^, 20%). The *k*_obs2_ (*k*_+α_) and the amplitude *A*_2_, did not show significant dependence on the varying [ATP]. The slow phase had a *k*_obs2_ of 63 ± 13 s^-1^ (Table 1). The *k*_obs1_ showed hyperbolic dependence on [ATP] with myosin S1 in the absence (control) ((Figure 8B B, green, filled circles) and in the presence of Mit (Figure 8B, pink, filled circles), but *k*_ob2_ is relatively independent of [MgATP] for control and Mit and saturates at 118 ± 79 s^-1^ and 79 ± 20 ^s-1^, respectively. The *k*_obs1_ for the other nine compounds showed similar hyperbolic dependence on [MgATP] that ranged between the control and that observed for Mit (Figure 8B).

**Table 1.** Kinetics parameters for the effects of compounds on ATP binding and isomerization in the actin-compound-S1 complex. Experimental conditions for stopped-flow experiments: F-buffer, 25°C, values are mean ± SEM for 4-7 replicates. ORIGIN fitting errors are reported for calculated equilibrium and rate constants. The effects for *K*_T_ and *k*_+2_ are categorized as: Mild (blue): < 2-fold; moderate (violet): 2-to 3-fold; severe (pink): >3-fold.

| Kinetics Parameter | Control | Car | Hon | Phn | Flp | Flu | Thi | Mef | Teg | Dan | Mit |
| --- | --- | --- | --- | --- | --- | --- | --- | --- | --- | --- | --- |
| **ATP binding & A.M.T → A~M.T isomerization** |  |  |  |  |  |  |  |  |  |  |  |
| $K_1k_{+2}$<br>($\mu\text{M}^{-1} \text{s}^{-1}$) | 3.83 ±<br>0.25 | 3.5 ±<br>0.3 | 4.14<br>± 0.3 | 6.4 ±<br>0.32 | 4.11<br>± 0.5 | 6.4 ±<br>0.3 | 4.9<br>±<br>0.4 | 5.1 ±<br>0.4 | 3.73<br>± 0.3 | 6.1 ±<br>0.8 | 6.2 ±<br>0.5 |
| $k_{+2}$ ($\text{s}^{-1}$) | 1621 ±<br>78 | 1196<br>± 78 | 1176<br>± 62 | 892<br>± 40 | 871 ±<br>39 | 843 ±<br>21 | 787<br>± 21 | 742<br>± 20 | 689 ±<br>20 | 649<br>± 53 | 635<br>± 29 |
| $K_T$ ($\mu\text{M}$) | 421 ±<br>34 | 342 ±<br>37 | 284 ±<br>27 | 139<br>± 9 | 212 ±<br>28 | 132 ±<br>7 | 161<br>± 14 | 146<br>± 12 | 185 ±<br>15 | 106 ±<br>16 | 102<br>± 10 |
| **AM → AM' isomerization** |  |  |  |  |  |  |  |  |  |  |  |
| $k_{+\alpha}$ ($\text{s}^{-1}$) | 63 ± 13 | 57 ±<br>11 | 59 ±<br>8 | 60 ±<br>12 | 51 ±<br>9 | 65 ±<br>11 | 54 ±<br>10 | 60 ±<br>9 | 42 ±<br>7 | 53 ±<br>8 | 59 ±<br>9 |
| $A_1$ | 0.81 ±<br>0.04 | 0.805<br>±<br>0.04 | 0.80<br>±<br>0.05 | 0.85<br>±<br>0.03 | 0.85<br>±<br>0.03 | 0.85<br>±0.03 | 0.86<br>±<br>0.02 | 0.81<br>±<br>0.04 | 0.83<br>±0.03 | 0.82<br>±<br>0.04 | 0.81<br>±<br>0.04 |
| $A_2$ | 0.19 ±<br>0.04 | 0.195<br>±<br>0.04 | 0.20<br>±<br>0.05 | 0.15<br>±<br>0.03 | 0.15<br>±<br>0.03 | 0.15<br>±<br>0.03 | 0.14<br>±<br>0.02 | 0.19<br>±<br>0.04 | 0.17<br>±<br>0.03 | 0.18<br>±<br>0.04 | 0.19<br>±<br>0.04 |
| $K_\alpha$<br>($A_1/A_2$) | 4.2 ±<br>0.8 | 4.11<br>± 0.8 | 3.87<br>± 1.0 | 5.67<br>± 1.2 | 5.76<br>± 1.0 | 5.7 ±<br>1.0 | 6.0<br>±<br>1.0 | 4.25<br>± 0.8 | 4.83<br>± 0.6 | 4.7<br>± 1.0 | 4.2<br>± 1.0 |

**Figure 8.**
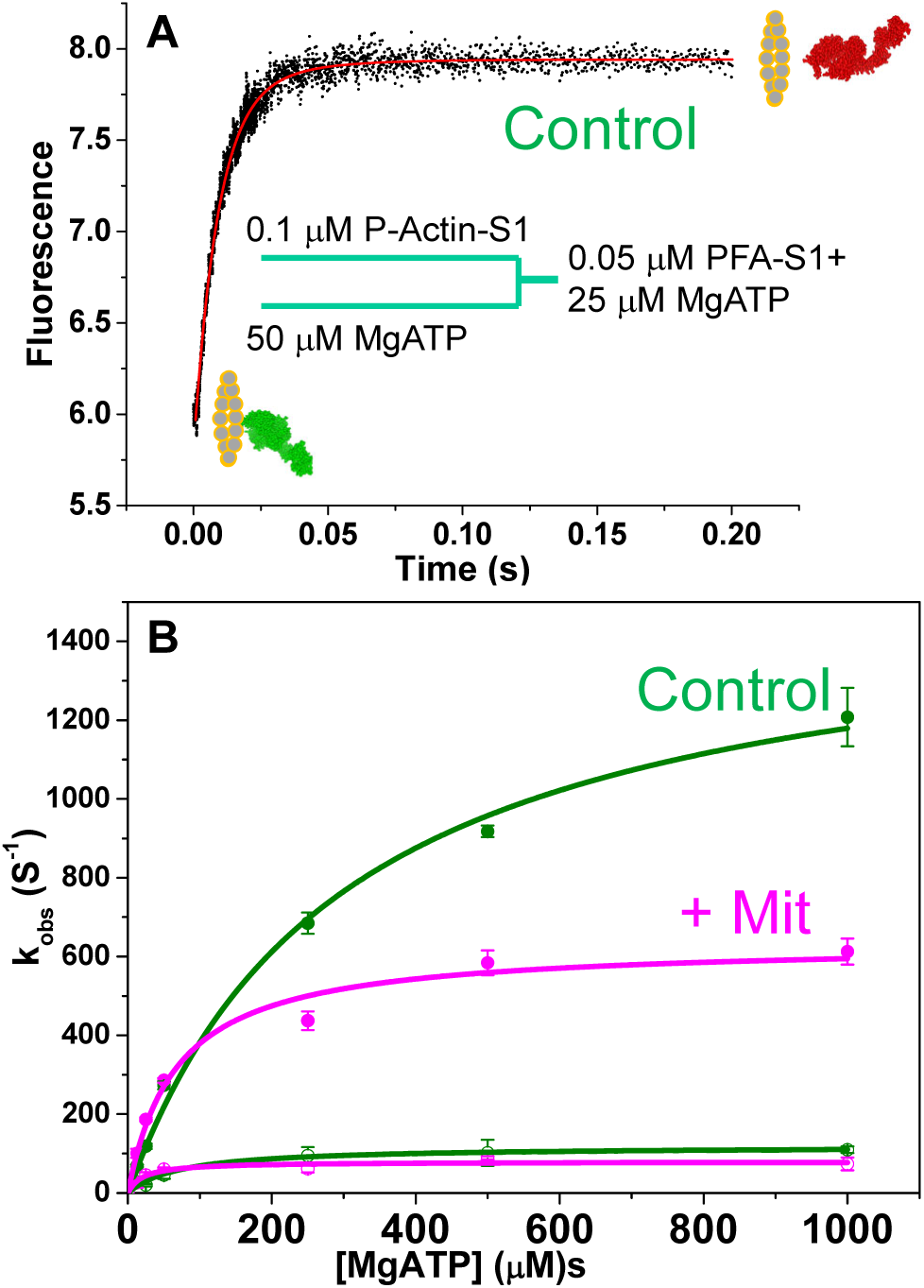
**A**: Transient kinetics of the dissociation of the complex of 0.05 μM PFA + 0.05 μM cardiac myosin S1 with 12.5μM MgATP. **B**: k_obs_ vs [MgATP] in the absence (green) and presence of 0.25 μM Mit (pink). *k*_obs1_ (filled circles) and *k*_obs2_ (open circles in corresponding colors)). Non-linear fits to a hyperbolic equation yields *K*_T_ or K_0.5_ (1/*K*_1_) and *k*_+2_ (**Table 1**).

The dissociation equilibrium constant *K*_T_ (1/*K*_1_) and rate constant *k*_+2_ are determined from these hyperbolic plots of *k*_obs1_ vs [MgATP] (Eq. 9) for ATP binding to the rigor actin-cardiac S1 complex (Figure 9A). For the control sample of actin and S1, we determined *K*_T_ and *k*_+2_ values of 421 ± 34 μM and 1621 ± 78 ^s-1^ (Table 1), respectively, which are consistent with that observed previously for cardiac β-S1 where K_T_ and *k*_+2_ were 328 ± 53 μM and 1543 ± 100 s^-1^ (2). All of the compounds induced decreases to varying extents in *K*_T_ and *k*_+2_ (Table 1, Figure 9B, C). Compared to the control values, we defined mild effects to be < 1.5-fold difference, moderate effects to be within 2-to 3-fold, and severe effects to be >3-fold. The dissociation equilibrium constant *K*_T_ is < 1.5-fold for Car and Hon compared to the control values (Figure 9B; Table 1). Mit and Dan induced the largest decrease (severe) in *K*_T_ (> 3.5-fold) (Figure 9B, pink, Figure 10). Similarly, Mef, Flu, and Phn also induced slightly greater than a 3-fold decrease (severe) in *K*_T_ (Figure 9B, pink, Figure 10). The other compounds (Flp, Thi, Teg) also significantly affected *K*_T_, decreasing it to varying extents to induce moderate effects (2 to 3-fold) (Figure 9B, violet, Table 1; Figure 10). The decreased dissociation equilibrium constants suggest that the “*affinity for ATP*” by myosin is increased in the presence of the compounds.

**Figure 9.**
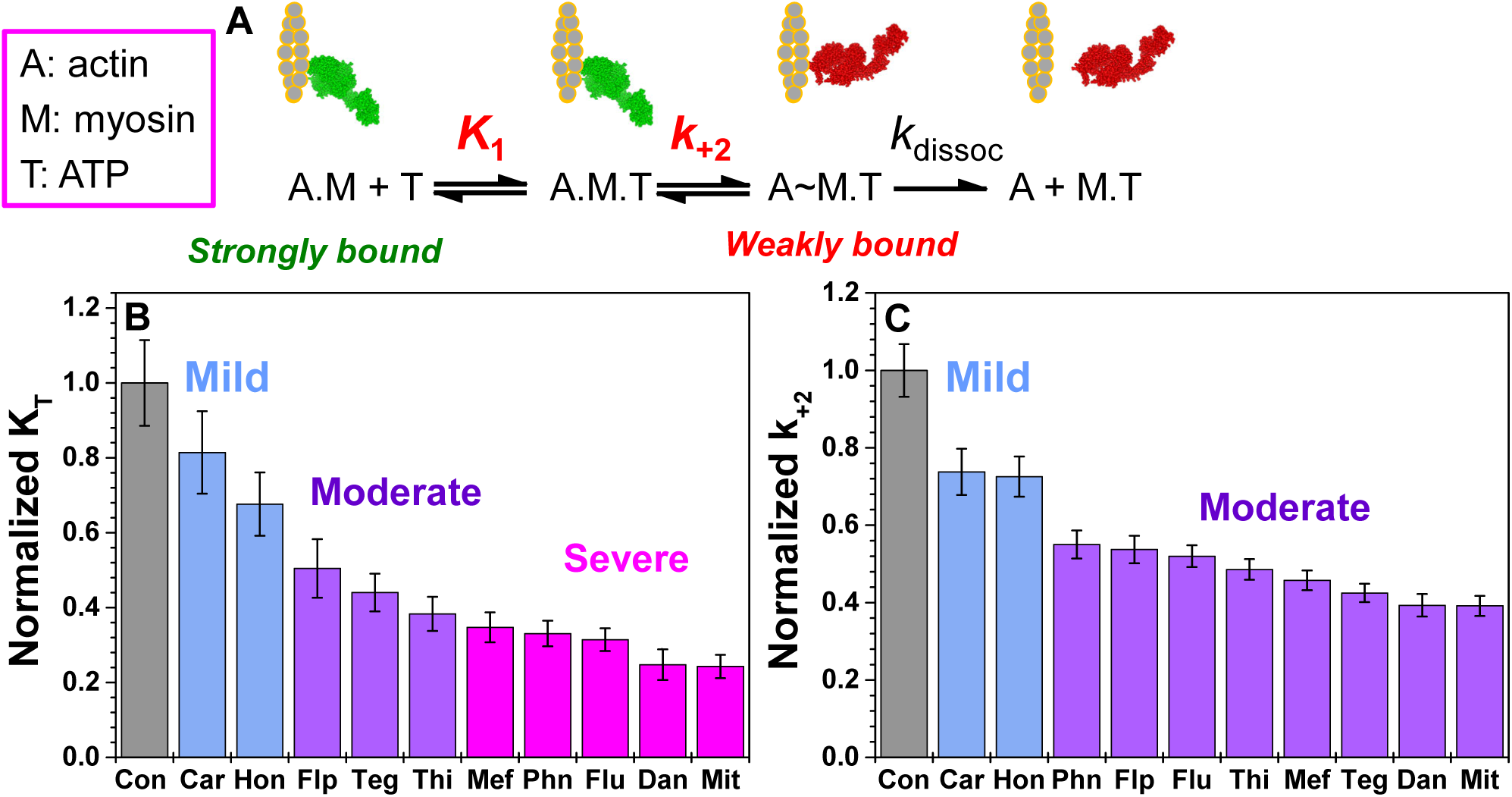
**A**: Kinetic scheme for myosin, showing binding and dissociation of ATP in the actin-myosin complex. **B**: Normalized equilibrium constant *K*_T_ (=1/*K*_1_), for cardiac S1 in the absence of the compound (grey) and in the presence of the compound. **C**: Normalized rate constant *k*_+2_, for cardiac S1 in the absence of the compound (grey) and in the presence of the compound. For control PFA and cardiac S1 in the absence of compounds (grey), the values are *K*_*T*_ = 421 ± 34 µM and *k*_*+*2_ = 1621 ± 78 s^-1^ (**Table 1**). The effects for *K*_T_ and *k*_+2_ are categorized as: Mild (blue): < 2-fold; moderate (violet): 2-to 3-fold; severe (pink): >3-fold.

**Figure 10.**
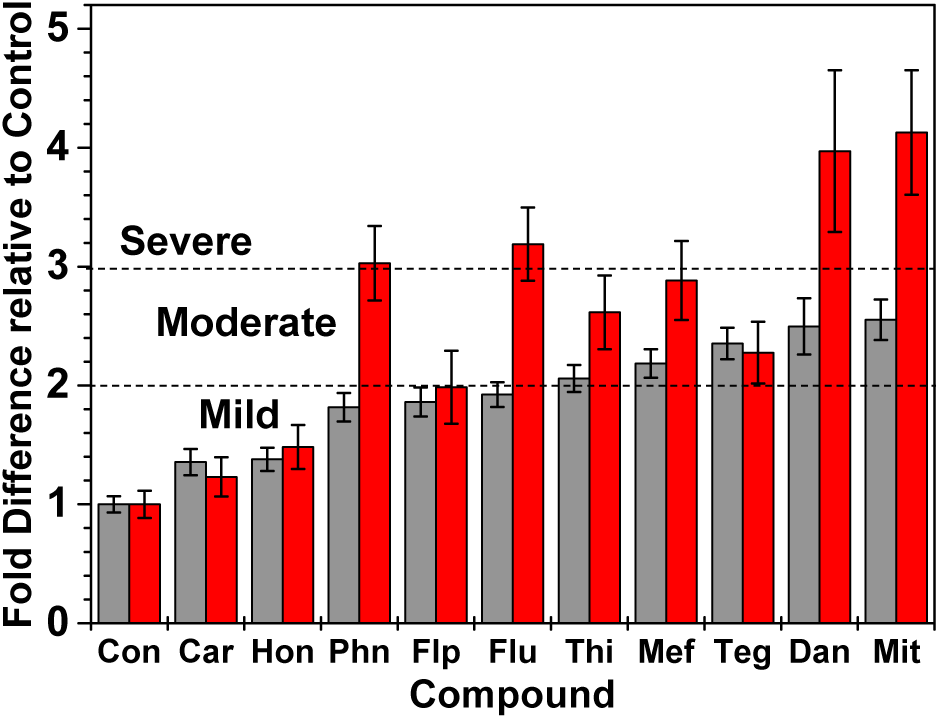
Comparison of the dissociation equilibrium constant, *K*_T_ (red), and the rate of isomerization of the collision complex to the ternary complex, *k*_+2_ (grey), for PFA and S1 in the absence (control) and presence of compounds. Dashed line shows 2-fold and 3 fold differences compared to control values. The effects are categorized as: Mild: < 2-fold; moderate: 2-to 3-fold; severe: >3-fold.

The rate constant, *k*_+2_, for the isomerization of the collision complex (A.M.T) to the ternary complex (A∼M.T) was determined from the hyperbolic fits of *k*_obs1_ vs [MgATP] (Eq. 9) in (shown in Figure 9). Dan and Mit showed the largest decrease in *k*_+2_ (> 2.5-fold) (Figure 9C, violet; Figure 10, Table 1) compared to the control sample (Figure 9C, grey). Flp, Phn and Flu also induced moderate decreases on *k*_+2_ (1.8-1.9 fold) (Figure 9B, violet, Figure 10). Thi, Mef and Teg also induced moderate decreases in *k*_+2_ (2-2.4 fold), while Car and Hon did not induce significant changes (<1.5 fold) in *k*_+2_ (Figure 9C, blue; Figure 10, Table 1).

We categorize the compound as severe if at least one parameter was affect >3-fold. Therefore, Mit, Dan, Mef, Flu, and Phn were determined to induce severe effects Flp, Thi, and Teg induces moderate effects, and Car and Hon induced mild effects (Figure 10).

The compounds did not significantly affect the average rate constant of the slow component, *k*_obs2_ (*k*_+α_), or the ratio of the amplitudes *A*_1_ and *A*_2_, or the equilibrium constant, *K*_α_, for the isomerization of actin.myosin from the closed (A.M) to open (A.M*’*) configuration (Scheme 2) to allow ATP binding to myosin (Table 1).

## Discussion

### Pyrene-Actin Fluorescence

Our steady-state fluorescence results show that the binding of cardiac S1 to F-actin linearly decreased the pyrene fluorescence up to a molar ratio of 1:1, as we previously showed (23). The inhibition of fluorescence is relieved by MgATP (Figure 2). PFA fluorescence is also sensitive to intermediate states during the weak-strong transition as evidenced by the sensitivity to nucleotide analogs that trap myosin in different bound nucleotide states such as ATPγS, ADP.VO_4_, PP_i_ (29,30); ADP.BeF_(x)_, ADP.AlF_4_, ADP.VO_4_ (31); and PP_i_ (27). Therefore, intermediate PFA fluorescence decrease could be reflective of either trapping nucleotide states in myosin (slower *k*_+2_ or decreased *k*_max_) or incomplete or partial dissociation of myosin from actin.

The addition of the compounds to PFA decreased the steady-state pyrene fluorescence to varying extents (Figure 4). The dissociation equilibrium constant K_d_ was determined from the change in PFA fluorescence as a function of [compound]. Mit, Flp, Dan, Thi, Mef, and Flu had a K_d_ that ranged from ∼ 5-50 μM, compared to the compounds that likely had a K_d_ > 100 μM. The compounds that showed tighter binding to actin (K_d_ < 50 μM) (Figure 4) are the compounds that induced the largest dysfunction in the actin-myosin interaction (Figure 9, Table 1). Our observation that the compounds directly affect the pyrene fluorescence of F-actin suggests that the binding of the compounds to actin induces a conformational change that exposes the pyrene moiety to the solvent, rendering it susceptible to collisions with other molecules that decrease its fluorescence. Our observation that MgATP relieves the fluorescence inhibition for some compounds (Mit) suggests that those compounds are bound near the nucleotide pocket in F-actin, which induces a conformational change that perturbs the bound compound, therefore, relieving the conformational change imposed at C374 in actin. In fact, Mit was identified in another HTS screen, which showed that Mit prevents GTPase binding to RhoGTApases (RhoA/RacA/Cdc42), and that Mit binds close to the GTP active site (32). It is unknown whether the compounds completely dissociates from F-actin in the presence of ATP.

### Rigor binding of Actin and S1

Some of the compounds (Dan, Mit, and Mef) inhibited the expected S1-induced decrease in steady-state PFA fluorescence and others inhibited it by even smaller extents (Flu, Flp, and Teg) (Figure 6A), suggesting that the presence of the compounds decreased the affinity of S1 for actin in rigor. Three other compounds (Phn, Thi, and Car) induced a larger pyrene fluorescence decrease than expected (Figure 6A), suggesting that these compounds induced stronger binding of S1 to actin. Hon did not affect the binding of S1 to actin.

We determined the rigor binding of actin to β-S1 by varying the [actin] in the range of two-to 12-fold in excess of S1 during the stopped-flow fluorescence transient kinetic experiments. The equilibrium dissociation constant, *K*_A_, for β-S1 was 15 ± 2 nM (Figure 7). This is consistent with previous reports of cardiac S1 from various species; for tissue-purified bovine cardiac β-S1 *K*_A_ was 6.3 nM (33), for tissue-purified human cardiac (HC) β-S1 *K*_A_ was 8 ± 2 nM or 10 ± 1.8 nM (2), and for bovine masseter β-S1 *K*_A_ was 7 nM (34). The dissociation equilibrium constant, *K*A, for actin binding to S1 in the presence of Mit and Dan, were 3-fold and 3.8-fold higher, respectively (Figure 7), suggesting that these compounds decreased the binding or association of PFA.compound complex with β-S1, which is consistent with the steady-state fluorescence results (Figure 6B) that showed incomplete S1 decrease of pyrene fluorescence in the presence of the compounds. Flu induced a smaller inhibition of the S1-induced decrease in the pyrene fluorescence, and the measured *K*_A_ was near normal values.

### ATP-induced dissociation of actin-S1 complex

Steady-state fluorescence showed that the compounds (except Car) inhibited the full restoration of PFA fluorescence (Figure 6B) due to ATP-induced dissociation of the actin-S1 complex, suggesting that the compounds induced conformational changes in actin that prevented the weakly-bound state of S1 to actin possibly by perturbing the myosin ATPase cycle. This is confirmed by the transient-state fluorescence measurements showed that the compounds decreased the equilibrium constant, *K*_T_, for ATP affinity for S1, and also decreased the subsequent rate constant, *k*_+2_, for the isomerization of the collision complex (A.M.T) to the ternary complex (A∼M.T) (Figure 9), therefore disrupting the myosin ATPase cycle and preventing complete and normal progression of the ATPase cycle. It is likely that the fully dissociated state of myosin is inhibited, which will also inhibit the hydrolysis of ATP in the unbound myosin and thus prevent the re-priming of the myosin head to allow for ADP release in subsequent steps of the ATPase cycle. Therefore, we hypothesize that the ATP is essentially “trapped” in the active site. Though the results were varied for the compounds, we categorized the compounds into three groups based on the extent of the fold difference compared to control values: mild (Car and Hon), moderate (Flp, Thi, and Teg), and severe (Mef, Phn, Flu, Mit, and Dan) (Figure 10).

### Mechanism of compound inhibition

Because these compounds decrease both the equilibrium dissociation constant *K*_T_ (*K*_m_), and the rate constant *k*_+2_ (*k*_max_), their mode of action is uncompetitive inhibition, in which the inhibitor binds to and stabilizes the enzyme-substrate complex allosterically (35). These allosteric effects are likely, as actin is known to act in a cooperative manner to affect either other actin monomers or myosin (36). It was previously shown that the myosin steady-state ATPase activity is unaffected by the presence of the compounds in the absence of actin (14), so inhibition is through conformational changes in actin that affect the M.ATP and M.ADP.P complex while bound to actin. This supports our hypothesis that the compounds trap ATP in the nucleotide site of myosin.

### Relationship to Other Work

Previous work in this laboratory showed that FRET from a fluorescent donor on the C-terminus of actin to an acceptor on the N-terminal extension of the cardiac myosin essential light chain (A1-ELC) is sensitive to disease-causing mutations in myosin (22). A subsequent study showed that a 12-amino-acid peptide derived from this N-terminal extension, also labeled with an acceptor, is sufficient to detect this interaction for the purposes of HTS (14,15). This led to the discovery of the ten actin-binding compounds in the present study, each of which causes a decrease in FRET efficiency, indicating that these compounds weaken the binding of cardiac myosin-derived peptide to actin (14). This was confirmed by the observation that actin-activated ATPase activity was decreased by several of the compounds (14). In a subsequent study, the same authors showed that several of these compounds showed significantly different effects on the Ca^2+^-dependence of steady-state ATPase activities of cardiac and skeletal myofibrils (16). This motivated the present study, in which we have used PFA and transient kinetics to define more clearly the mechanisms of action of these compounds, focusing on cardiac actin-myosin interaction.

### Structural mechanism of action of the compounds

Holmes and co-workers (37) showed that myosin binds initially in a weak stereospecific interaction with actin, in which the myosin nucleotide-binding cleft is closed, while strong binding of S1 to actin opens the nucleotide-binding pocket to allow binding of the nucleotide. Our results show that some of the compounds (Mit, Dan, Mef, Flu, Flp, and Teg) decrease the effect of S1 on PFA fluorescence, suggesting weaker actin binding, while other compounds (Phn, Thi, and Car) increase the effect of S1, suggesting stronger actin binding.

The compounds that decreased strong-binding of S1 to actin also significantly decreased the dissociation equilibrium constant *K*_T_ for ATP binding to actin-S1, suggesting stronger binding of ATP to S1 in the S1.actin complex. These compounds also decreased the rate of isomerization from the collision complex (A.M.T) to the ternary complex (A∼M.T), thus possibly disrupting the W→S transition myosin undergoes during the power stroke. It is likely that structural changes induced by these compounds in actin affects myosin structure, trapping ATP in its active site, inhibiting the “priming” of myosin for actin reattachment and normal completion of the ATPase cycle.

### Potential therapeutic applications

Two compounds that we categorized as severe and moderate (Mit, Teg) have been shown to induce heart failure; Thi and Flp (moderate effects in the present study) were removed from clinical use because they caused liver toxicity. Although most of the compounds in the present study have been used to treat non-contractile dysfunction in humans (as anti-cancer, anti-psychotic, and anti-malarial agents), it is possible that some of them, or derivatives of them, have therapeutic potential for cardiomyopathy where attenuation of contractility is beneficial, as in a hypercontractile state caused by HCM mutations in myosin or actin (38,39). Regulation of striated muscle contractility is not only thin-filament dependent, but also thick-filament dependent, whereby the myosin binding affinity to actin is attenuated via phosphorylation of the light chains (40) and myosin-binding protein C (41,42), changes in flexibility and conformation (LC domain (43), converter region (44), actin binding region (3,45), and small molecules (Mava and OM) (39) that affect the myosin ATPase cycle. In the present study, we have shown that the compounds increase ATP affinity and decrease the rate constant for the subsequent isomerization steps in ternary complex formation (and hence the release of S1 from actin), which is likely to produce actin-bound non-force producing heads, resulting in decreased ATPase activity. It is possible that these actin-binding compounds may prove to be useful in decreasing hypercontractility in the HCM disease, caused by either actin or myosin mutations, by attenuating the binding affinities of myosin for actin.

### Conclusions

Starting from compounds previously identified from high-throughput screening, we have shown that pyrene-labeled actin is sensitive to conformational changes that these compounds induce in F-actin, and that these conformational changes affect the strong-binding and weak-binding states of myosin to actin. Some of these compounds prevent cardiac myosin S1 from decreasing pyrene-actin steady-state fluorescence in the absence of ATP, implying that they decreased the fraction of strongly bound myosin in rigor. Many of the compounds prevent full restoration of steady-state fluorescence from actin-S1 complex in ATP, implying that they either decrease the fraction of nucleotide-bound myosin, or that the actin-attached states are more populated in the presence of ATP.

Transient kinetics shows that the affinity for actin binding to S1 is weaker (increased equilibrium constant, *K*_A_) in the presence of Mit and Dan. In addition, the binding affinity of ATP for myosin in the rigor actin-myosin complex is increased for most of the compounds (possibly enhancing the formation of the collision complex, A.M.T), which then induces a subsequent decrease in the rate constant for the isomerization of the collision complex to the ternary complex, A∼M.T. Most of the other compounds show a large decrease in *K*_T_, but a smaller decrease in *k*_+2,_ thus revealing enhanced affinity for ATP in the actin-myosin complex and decreased rate of the subsequent isomerization step that results in impaired actin-activated myosin ATPase activity. Therefore, the present study shows that the early steps of the cardiac myosin ATPase kinetics cycle is affected by myosin structural changes induced by compounds binding to actin. To get further insight into the mechanisms of action of these compounds, further studies are needed to resolve the kinetics of ADP release (rate limiting step for β-cardiac myosin), phosphate release, the powerstroke, and contractility of skeletal and cardiac muscle fibers. Most importantly, this research also sets the stage for studying the mechanism of action of compounds discovered from future HTS studies, to detect new actin-binding compounds.

## Experimental procedures

### Preparations and Assays

Acetone powder was prepared from young female New Zealand White rabbit skeletal muscle (46). Rabbit skeletal F-actin was prepared as described previously from acetone powder (47). Bovine cardiac β-myosin and α-chymotryptic S1 were prepared from bovine left ventricular muscle (Pelfreez Biologicals, MO) (9,48) and stored in sterile 150 mM sucrose at -80°C until needed. The actin-activated myosin S1 steady-state MgATPase activity was measured using the NADH-coupled assay at 25°C using 15 μM phalloidin-stabilized F-actin and 0.1 μM cardiac β-S1 in F-Mg buffer (10mM Tris (pH 7.5), 3mM MgCl_2_) (22,23). ATPase activity was reported as the mean ± SEM s^-1^ (n = 4).

The ten compounds were obtained from Target Molecule Corp.(Boston, MA) and stored at - 20°C as 10 mM stock solutions in DMSO until use. Omecamtiv mecarbil (OM) and mavacamten (Mava) were obtained from Selleck Chemicals (Houston, TX), and stored as 10 mM stock solutions at -20°C until use. Sodium vanadate and potassium pyrophosphate (PP_i_) were obtained from Sigma-Aldrich (St. Louis, MO), and stock solutions of 200 mM and 1M, respectively, were prepared as described previously (26).

### Labeling F-Actin

F-actin was labeled with N-(1-pyrene) iodoacetamide (PIA) (Invitrogen, Thermo Fisher Scientific, Waltham, MA) essentially as described previously (17,20). Briefly, F-actin (1mg/ml) in F-Mg buffer, was mixed with PIA at a molar ratio of 1:10 and left overnight at room temperature in the dark. 10 mM DTT was used to stop the reaction, and the labeled F-actin was sedimented at 80K for 45 min at 4C. The pellet was resuspended with a dounce homogenizer in G-buffer (5 mM Tris (pH 7.5), 0.5 mM ATP, 0.2 mM MgCl_2_), centrifuged for 10 min at 70K to remove the PIA precipitate. The supernatant was repolymerized with 3 mM MgCl_2_, and the labeled F-actin was collected in the pellet after centrifugation at 80K for 45 min, then resuspended with a dounce homogenizer with F-Mg buffer. The pyrene concentration on F-actin was determined by absorbance (344nm) and extinction coefficient of 2.2 x104 M^-1^ cm^-1^. The labeled actin concentration was determined by Bradford assay (Biorad). A typical dye:protein ratio of 0.9-0.95 was obtained for pyrene-labeled actin. The pyrene-F-actin (PFA) was stabilized with phalloidin (Sigma-Aldrich, St. Louis, MO) at a 1:1 molar ratio, stored on ice, and used within a week.

### Steady-State Fluorescence Spectroscopy

Fluorescence spectra of 2 μM PFA in F-Mg buffer were acquired with a Cary spectrophotometer. The sample was placed in a cuvette in a temperature-controlled holder, and excited with a Xenon flash lamp at 365 nm with 5nm slit width. Emission fluorescence intensity was acquired and averaged for 30 sec at 407 nm with a 10 nm slit width. A final concentration of 2 μM PFA in F-actin buffer was used. Fluorescence intensity (F) was corrected for each compound’s emission at 407nm. Control experiments with the compounds in the absence of PFA showed minimal fluorescence emission (<1%) at 407 nm. The fractional fluorescence (*f*_f_) due to S1 binding to actin was determined by normalization relative to the PFA fluorescence, as described in Eq. 1:

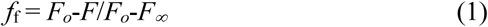

*F*_*o*_ is the fluorescence in the absence of S1, *F*_*∞*_ is the fluorescence at infinite [S1], and *F* is the fluorescence in the presence of S1 and/or compound.

The binding of a compound to PFA was determined from the dose-dependent change in PFA fluorescence in the range of 0-150μM compound concentration [C]. Control experiments with 1-1.5% DMSO showed negligible effect on PFA fluorescence. The fraction of compound bound (*f*_b_) was determined for the compounds that showed hyperbolic dependence of PFA fluorescence (*F*_o_) as a function of [C] using Eq. 2. The dissociation equilibrium constant (*K*_d_) for the compound binding to PFA was determined from hyperbolic fits of plots of *f*_b_ vs [C] that plateau at *f*_max_ (Eq. 3).

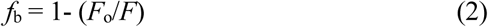

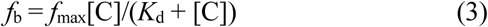

### Rigor binding of S1 to F-Actin-Compound complex

To determine whether a compound affected the rigor binding of S1 to F-Actin, the expected decrease in SS fluorescence intensity (F_calc_) was determined (Eq. 4). To ensure that a negative fluorescence intensity was not induced by the additive effect of the compound and S1, a ratio of 0.7:1 of S1:actin was used in the experiment. *F*_calc_ was determined from normalized fluorescence intensity:

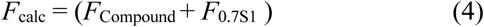

### Transient-state Fluorescence Spectroscopy

Transient kinetics experiments to detect the total fluorescence intensity were performed using an Applied Photophysics stopped-flow spectrophotometer that is capable of single-mix and sequential-mix experiments with temperature control (23). All experiments were performed at 25°C with filtered and degassed solutions. The instrument dead-time for single-mix experiments was 1.3 ms. Samples were excited at 365 nm, and pyrene emission fluorescence was acquired with a 2 mm path length and 400 nm long-pass filter.

### Analysis of stopped-flow fluorescence data

Fluorescence decays from the transient kinetics measurements were fitted to single (Eq. 5) and double exponential (Eq. 6) equations. The best fit was evaluated by the lowest χ^2^ value of the fit compared to the experimental data.

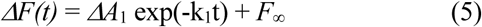

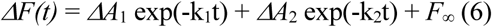

Δ*F* is the change in PFA fluorescence, *F*_*∞*_ is the fluorescence at infinite [substrate], and *A* and *k* are the observed amplitude and rate constant of the fluorescence change, respectively.

### Rigor binding of S1 to Actin

As PFA fluorescence is very sensitive to rigor binding of myosin, we determined the effect of each compound on the binding of S1 to actin. The rate of binding, *k*_+A_ was determined by varying the PFA concentration in the presence of S1. To remove any effects of ATP or ADP, the mixed actin-S1 samples were treated with 3 units of apyrase per mL for five minutes at room temperature before fluorescence acquisition. Addition of apyrase removed any contaminating nucleotide, and rendered the transient decays of the control experiments with cardiac S1 only as single exponentials (17,28). The linear plot of *k*_obs_ vs [PFA] yielded a slope that described the second-order rate constant (*k*_+A_) of S1 binding to actin, while the intercept described the rate constant rate of S1 dissociation (*k*_-A_) (Scheme 1, Eq. 7). The equilibrium constant for dissociation, *K*_A_, was determined from the ratio of the rate constants *k*-_A_/*k*_+A_ (Scheme 1, Eq. 8).

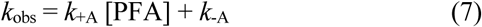

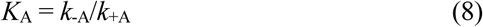

### Dissociation of the rigor actin-S1 complex with MgATP

The detachment of S1 from actin involves the binding of ATP (T) to the open configuration of A.M (Scheme 2), which is controlled by the diffusion-limited association equilibrium constant *K*_1_, in the formation of the collision complex (A.M.T), which is followed by the rate limiting (*k*+2) isomerization to A∼M.T, in which actin is weakly bound and ATP is more strongly bound to myosin, (Scheme 2) (17). This is rapidly followed by dissociation of myosin S1 from actin at a rate of *k*dissoc.

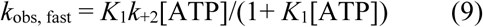

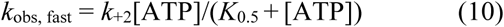

*K*_1_ was determined by mixing the rigor complex of PFA.S1 (A.M) with varying concentrations of MgATP in the absence and presence of the compounds. For cardiac β-myosin S1, the transient curves were fitted to rate constants, *k*_obs, fast_ and *k*_obs_,_slow_, and with corresponding amplitudes *A*_1_ and *A*_2_. At low concentrations of ATP (*K*_1_.MgATP << 1), the plot of the observed rate constant (*k*_obs_,_fast_) vs [MgATP] shows a linear dependence that is typical of a pseudo second-order reaction, from which the slope is the apparent second-order rate constant, *K*_1_*k*_+2_ (Eq. 9). However, at higher concentrations of MgATP, the relationship becomes hyperbolic (as described in Eq. 10) in which a plot of *k*_obs_ vs [Mg.ATP] plateaus at *k*_max_, which corresponds to rate constant *k*_+2_. The dissociation equilibrium constant of the actin-S1complex *K*_0.5*_(1/*K*_1_) = *K*_T_ is determined from *k*_+2_/*K*_1_*k*_+2_, and is the ATP concentration at half saturation that describes the binding “affinity of ATP” (*K*_T_) for the actin-myosin complex.

## Data Availability

All data described are contained within the article.

## Acknowledgments

Spectroscopic instrumentation was provided by the Biophysical Technology Center (BTC) at UMN. We thank Dr. John D. Lipscomb at UMN for use of the stopped-flow fluorescence instrument, and Dr. Rahul Banerjee and Dr. Melanie Rogers for technical assistance with that instrument and for helpful discussions on kinetics experiments and data analysis.

## Funding and Additional Information

This work was supported in part by NIH grants to DDT (R01AR032961 and R37AG26160) and to OR (R01AR052360).

---

**Scheme 1.**
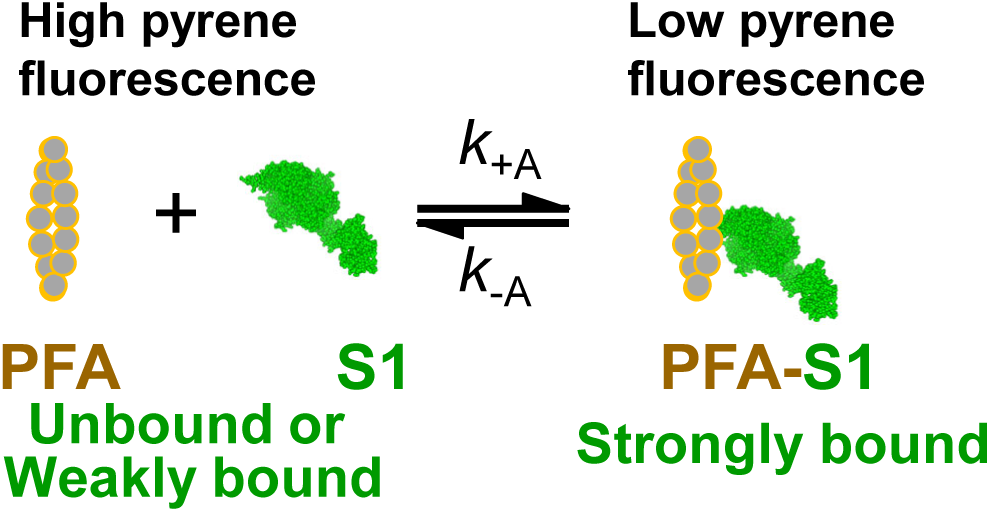
Actin binding to and dissociation from myosin S1.

**Scheme 2.**
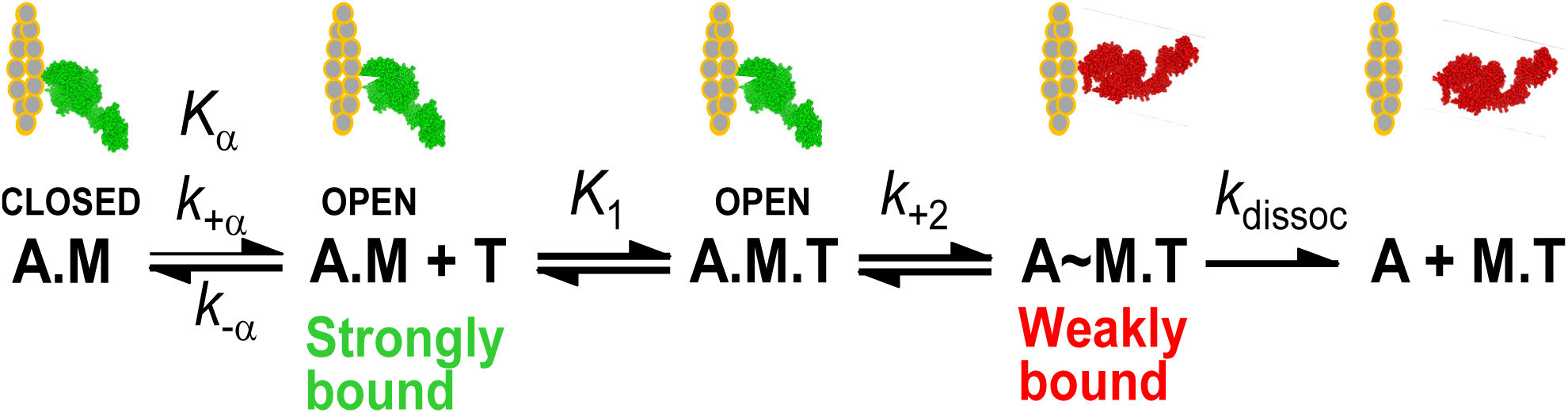
ATP-induced dissociation of actin-myosin complex.

## References

1. Lee, S. H., and Dominguez, R. (2010) Regulation of actin cytoskeleton dynamics in cells. Mol Cells 29, 311–325

2. Vera, C. D., Johnson, C. A., Walklate, J., Adhikari, A., Svicevic, M., Mijailovich, S. M., Combs, A. C., Langer, S. J., Ruppel, K. M., Spudich, J. A., Geeves, M. A., and Leinwand, L. A. (2019) Myosin motor domains carrying mutations implicated in early or late onset hypertrophic cardiomyopathy have similar properties. J Biol Chem 294, 17451–17462

3. Thomas, D. D., Prochniewicz, E., and Roopnarine, O. (2002) Changes in actin and myosin structural dynamics due to their weak and strong interactions. Results Probl Cell Differ 36, 7–19

4. Marston, S. (2018) The Molecular Mechanisms of Mutations in Actin and Myosin that Cause Inherited Myopathy. Int J Mol Sci 19

5. Despond, E. A., and Dawson, J. F. (2018) Classifying Cardiac Actin Mutations Associated With Hypertrophic Cardiomyopathy. Front Physiol 9, 405

6. Malik, F. I., Hartman, J. J., Elias, K. A., Morgan, B. P., Rodriguez, H., Brejc, K., Anderson, R. L., Sueoka, S. H., Lee, K. H., Finer, J. T., Sakowicz, R., Baliga, R., Cox, D. R., Garard, M., Godinez, G., Kawas, R., Kraynack, E., Lenzi, D., Lu, P. P., Muci, A., Niu, C., Qian, X., Pierce, D. W., Pokrovskii, M., Suehiro, I., Sylvester, S., Tochimoto, T., Valdez, C., Wang, W., Katori, T., Kass, D. A., Shen, Y. T., Vatner, S. F., and Morgans, D. J. (2011) Cardiac myosin activation: a potential therapeutic approach for systolic heart failure. Science 331, 1439–1443

7. Mamidi, R., Li, J., Gresham, K. S., Verma, S., Doh, C. Y., Li, A., Lal, S., Dos Remedios, C. G., and Stelzer, J. E. (2017) Dose-Dependent Effects of the Myosin Activator Omecamtiv Mecarbil on Cross-Bridge Behavior and Force Generation in Failing Human Myocardium. Circ Heart Fail 10

8. Green, E. M., Wakimoto, H., Anderson, R. L., Evanchik, M. J., Gorham, J. M., Harrison, B. C., Henze, M., Kawas, R., Oslob, J. D., Rodriguez, H. M., Song, Y., Wan, W., Leinwand, L. A., Spudich, J. A., McDowell, R. S., Seidman, J. G., and Seidman, C. E. (2016) A small-molecule inhibitor of sarcomere contractility suppresses hypertrophic cardiomyopathy in mice. Science 351, 617–621

9. Rohde, J. A., Thomas, D. D., and Muretta, J. M. (2017) Heart failure drug changes the mechanoenzymology of the cardiac myosin powerstroke. Proc Natl Acad Sci U S A 114, E1796–E1804

10. Nanasi, P., Jr., Komaromi, I., and Almassy, J. (2018) Perspectives of a myosin motor activator agent with increased selectivity. Can J Physiol Pharmacol 96, 676–680

11. Nanasi, P., Komaromi, I., Gaburjakova, M., and Almassy, J. (2018) Omecamtiv Mecarbil: A Myosin Motor Activator Agent with Promising Clinical Performance and New in vitro Results. Curr Med Chem 25, 1720–1728

12. Fumagalli, C., De Gregorio, M. G., Zampieri, M., Fedele, E., Tomberli, A., Chiriatti, C., Marchi, A., and Olivotto, I. (2020) Targeted Medical Therapies for Hypertrophic Cardiomyopathy. Curr Cardiol Rep 22, 10

13. Sparrow, A. J., Watkins, H., Daniels, M. J., Redwood, C., and Robinson, P. (2020) Mavacamten rescues increased myofilament calcium sensitivity and dysregulation of Ca(2+) flux caused by thin filament hypertrophic cardiomyopathy mutations. Am J Physiol Heart Circ Physiol 318, H715–H722

14. Guhathakurta, P., Prochniewicz, E., Grant, B. D., Peterson, K. C., and Thomas, D. D. (2018) High-throughput screen, using time-resolved FRET, yields actin-binding compounds that modulate actin-myosin structure and function. J Biol Chem 293, 12288–12298

15. Guhathakurta, P., Prochniewicz, E., and Thomas, D. D. (2018) Actin-Myosin Interaction: Structure, Function and Drug Discovery. Int J Mol Sci 19

16. Guhathakurta, P., Phung, L. A., Prochniewicz, E., Lichtenberger, S., Wilson, A., and Thomas, D. D. (2020) Actin-binding compounds, previously discovered by FRET-based high-throughput screening, differentially affect skeletal and cardiac muscle. J Biol Chem

17. Adamek, N., and Geeves, M. A. (2014) Use of pyrene-labelled actin to probe actin-myosin interactions: kinetic and equilibrium studies. Exp Suppl 105, 87–104

18. Onishi, H., Mikhailenko, S. V., and Morales, M. F. (2006) Toward understanding actin activation of myosin ATPase: the role of myosin surface loops. Proc Natl Acad Sci U S A 103, 6136–6141

19. O’Donoghue, S. I., Miki, M., and dos Remedios, C. G. (1992) Removing the two C-terminal residues of actin affects the filament structure. Arch Biochem Biophys 293, 110–116

20. Criddle, A. H., Geeves, M. A., and Jeffries, T. (1985) The use of actin labelled with N-(1-pyrenyl)iodoacetamide to study the interaction of actin with myosin subfragments and troponin/tropomyosin. Biochem J 232, 343–349

21. Kouyama, T., and Mihashi, K. (1981) Fluorimetry study of N-(1-pyrenyl)iodoacetamide-labelled F-actin. Local structural change of actin protomer both on polymerization and on binding of heavy meromyosin. Eur J Biochem 114, 33–38

22. Guhathakurta, P., Prochniewicz, E., Roopnarine, O., Rohde, J. A., and Thomas, D. D. (2017) A Cardiomyopathy Mutation in the Myosin Essential Light Chain Alters Actomyosin Structure. Biophys J 113, 91–100

23. Rohde, J. A., Roopnarine, O., Thomas, D. D., and Muretta, J. M. (2018) Mavacamten stabilizes an autoinhibited state of two-headed cardiac myosin. Proc Natl Acad Sci U S A 115, E7486–E7494

24. Goodno, C. C. (1979) Inhibition of myosin ATPase by vanadate ion. Proc Natl Acad Sci U S A 76, 2620–2624

25. Goodno, C. C., and Taylor, E. W. (1982) Inhibition of actomyosin ATPase by vanadate. Proc Natl Acad Sci U S A 79, 21–25

26. Roopnarine, O., and Thomas, D. D. (1996) Orientation of intermediate nucleotide states of indane dione spin-labeled myosin heads in muscle fibers. Biophys J 70, 2795–2806

27. Taylor, E. W. (1991) Kinetic studies on the association and dissociation of myosin subfragment 1 and actin. J Biol Chem 266, 294–302

28. De La Cruz, E. M., and Ostap, E. M. (2009) Kinetic and equilibrium analysis of the myosin ATPase. Methods Enzymol 455, 157–192

29. Geeves, M. A., and Jeffries, T. E. (1988) The effect of nucleotide upon a specific isomerization of actomyosin subfragment 1. Biochem J 256, 41–46

30. Geeves, M. A., Jeffries, T. E., and Millar, N. C. (1986) ATP-induced dissociation of rabbit skeletal actomyosin subfragment 1. Characterization of an isomerization of the ternary acto-S1-ATP complex. Biochemistry 25, 8454–8458

31. Xu, J., and Root, D. D. (2000) Conformational selection during weak binding at the actin and myosin interface. Biophys J 79, 1498–1510

32. Bidaud-Meynard, A., Arma, D., Taouji, S., Laguerre, M., Dessolin, J., Rosenbaum, J., Chevet, E., and Moreau, V. (2013) A novel small-molecule screening strategy identifies mitoxantrone as a RhoGTPase inhibitor. Biochem J 450, 55–62

33. Siemankowski, R. F., and White, H. D. (1984) Kinetics of the interaction between actin, ADP, and cardiac myosin-S1. J Biol Chem 259, 5045–5053

34. Bloemink, M. J., Adamek, N., Reggiani, C., and Geeves, M. A. (2007) Kinetic analysis of the slow skeletal myosin MHC-1 isoform from bovine masseter muscle. J Mol Biol 373, 1184–1197

35. Ring, B., Wrighton, S. A., and Mohutsky, M. (2014) Reversible mechanisms of enzyme inhibition and resulting clinical significance. Methods Mol Biol 1113, 37–56

36. Prochniewicz, E., and Thomas, D. D. (1997) Perturbations of functional interactions with myosin induce long-range allosteric and cooperative structural changes in actin. Biochemistry 36, 12845–12853

37. Holmes, K. C., Angert, I., Kull, F. J., Jahn, W., and Schroder, R. R. (2003) Electron cryo-microscopy shows how strong binding of myosin to actin releases nucleotide. Nature 425, 423–427

38. Spudich, J. A. (2019) Three perspectives on the molecular basis of hypercontractility caused by hypertrophic cardiomyopathy mutations. Pflugers Arch 471, 701–717

39. Yotti, R., Seidman, C. E., and Seidman, J. G. (2019) Advances in the Genetic Basis and Pathogenesis of Sarcomere Cardiomyopathies. Annu Rev Genomics Hum Genet 20, 129–153

40. Gordon, A. M., Homsher, E., and Regnier, M. (2000) Regulation of contraction in striated muscle. Physiol Rev 80, 853–924

41. Colson, B. A., Thompson, A. R., Espinoza-Fonseca, L. M., and Thomas, D. D. (2016) Site-directed spectroscopy of cardiac myosin-binding protein C reveals effects of phosphorylation on protein structural dynamics. Proc Natl Acad Sci U S A 113, 3233–3238

42. Wang, L., Geist, J., Grogan, A., Hu, L. R., and Kontrogianni-Konstantopoulos, A. (2018) Thick Filament Protein Network, Functions, and Disease Association. Compr Physiol 8, 631–709

43. Savich, Y., Binder, B. P., Thompson, A. R., and Thomas, D. D. (2019) Myosin lever arm orientation in muscle determined with high angular resolution using bifunctional spin labels. J Gen Physiol 151, 1007–1016

44. Gunther, L. K., Rohde, J. A., Tang, W., Walton, S. D., Unrath, W. C., Trivedi, D. V., Muretta, J. M., Thomas, D. D., and Yengo, C. M. (2019) Converter domain mutations in myosin alter structural kinetics and motor function. J Biol Chem 294, 1554–1567

45. Colson, B. A., Gruber, S. J., and Thomas, D. D. (2012) Structural dynamics of muscle protein phosphorylation. J Muscle Res Cell Motil 33, 419–429

46. Pardee, J. D., and Spudich, J. A. (1982) Purification of muscle actin. Methods Enzymol 85 Pt B, 164–181

47. Prochniewicz, E., Zhang, Q., Howard, E. C., and Thomas, D. D. (1996) Microsecond rotational dynamics of actin: spectroscopic detection and theoretical simulation. J Mol Biol 255, 446–457

48. Margossian, S. S., and Lowey, S. (1982) Preparation of myosin and its subfragments from rabbit skeletal muscle. Methods Enzymol 85 Pt B, 55–71

